# Sharing the Spotlight: Uncovering common attentional dynamics across species

**DOI:** 10.1101/2024.11.01.621490

**Authors:** Mina Glukhova, Alejandro Tlaie, Raul Muresan, Robert Taylor, Pierre-Antoine Ferracci, Katharine Shapcott, Olga Arne, Andrei Ciuparu, Martha N Havenith, Marieke L Schölvinck

**Affiliations:** Ernst Strüngmann Institute for Neuroscience in cooperation with the Max Planck Society, Frankfurt am Main, 60528, Germany; Transylvanian Institute of Neuroscience, 400157 Cluj-Napoca, Romania; STAR-UBB Institute, Babeş-Bolyai University, 400084 Cluj-Napoca, Romania

**Author notes:** These authors contributed equally (Martha N Havenith, Marieke L Schölvinck).

**Keywords:** Sustained attention, Cross-species, Naturalistic behaviour, Virtual Reality

## Abstract

Sustained attention is a key underlying process to many natural behaviours that are shared by multiple species. Yet the way it is commonly studied in a lab context precludes any meaningful cross-species comparisons. Here, we engaged mice, monkeys, and humans in the exact same, natural perceptual decision task in a VR environment. We captured their behaviour into several parameters along the speed/accuracy axes along which sustained attention is classically defined, and used HMMs to infer four attentional states. We show that the dynamics of these states, both in terms of their durations as well as the transitions between them, are much more similar between the species than might have been expected. Moreover, attentional state fluctuations seem to be internally generated. The task and analyses developed here represent a new approach to compare sustained attention across species in an objective, data-driven way.

## Introduction

Compared to insects and invertebrates, many mammals have a large repertoire of flexible behaviours, such as foraging for food, scanning the environment for potential dangers, and social interactions with conspecifics. For many of these behaviours, attention is a key guiding cognitive process. Without sustained attention, mice would be easily caught by predator birds; monkeys would not find enough food; and humans would not be able to read scientific papers.

In a neuroscientific context, sustained attention is commonly translated into attention towards a certain task, also known as task engagement. A state of high sustained attention manifests itself in certain physiological parameters, such as increased pupil size (1, 2) and increased locomotion (3). A state of high attention can typically not be sustained for extended periods of time (4); as anyone trying to stay focused on a difficult task for hours can testify, sustained attention fluctuates naturally (5, 6).

The hallmark of high sustained attention, however, is better task performance: animals or humans attending to a task are typically faster and less variable in their response times (7, 8) and have more correct responses (9, 10). Importantly, speed and accuracy often do not go together; the well-known speed-accuracy trade-off denotes the common phenomenon whereby participants either prioritize speed or accuracy as an attentional strategy in allocating limited processing resources (11). They can be biased to prioritize either strategy through reward incentives (12, 13), although reward prospect per se also enhances overall performance (14).

Given that it is such a fundamental building block of cognition, it seems reasonable to assume that dynamically fluctuating sustained attention is largely shared between the species. Yet it is difficult to meaningfully compare attentional fluctuations across species given that their behaviours are typically measured in highly divergent task contexts. Mice, for instance, are typically engaged in sensory discrimination tasks that involve strong motor components, such as repeatedly traversing an experimental chamber to reach different buttons, nose pokes locations or touch screens, or running on a treadmill and licking for reward. Sustained attention towards the task is then inferred from metrics such as the amount of movement unrelated to the task (10, 15) or from the amount of anticipatory licking (13, 16).

In non-human primate experiments, this form of embodied cognition is rare. They are typically highly overtrained on demanding (mostly) visual tasks, that require them to sit still and only engage in highly automatized responses in the form of small eye movements or button presses. The few studies investigating natural performance fluctuations in such a task context have found that sustained attention has a strong effect on task accuracy: it greatly improves the detection of subtle changes in a stimulus, but impairs the ability to notice when that stimulus changes dramatically (17). Attentional fluctuations in such a restricted primate task seem to be modest from trial to trial (18), sometimes interspersed with lapses of complete task disengagement (19).

In humans, sustained attention has been studied extensively using continuous performance tasks (CPTs); repetitive, boring tasks that require them to maintain their focus in order to respond to targets or inhibit responses to foils. For example, in the sustained attention to response task (SART), participants are required to respond on most trials, with rare “stop” trials serving as targets (20). CPTs have been used to expose the speed-accuracy tradeoff; prolonged performance leads to faster and more error-prone responding (21, 22). Lapses in sustained attention manifest as failures to withhold response (20), i.e. a drop in task accuracy.

Originally developed for humans, CPTs have been adapted to touchscreen versions, to be used in monkeys (23, 24) as well as rodents (25, 26). Findings show that performance on this task by rodents was highly translatable to human performance (25). Though trainable in monkeys and mice, CPTs elicit very unnatural and limited behaviour, and therefore it is unknown whether these findings generalise to other task paradigms - and how well they reflect attentional fluctuations that might occur spontaneously in the wild.

In order to study sustained attention in natural behaviour across species, we engaged mice, monkeys, and humans in the exact same perceptual decision task. To facilitate comparison of spontaneous attentional dynamics across species, we made the task as naturalistic as possible: Instead of an abstract mapping of e.g. button presses to artificial stimuli, stimuli resembling natural objects were displayed in a virtual reality (VR) environment and subjects ‘walked’ towards them. While walking (for mice by running on a spherical treadmill, for monkeys and humans by moving a trackball with their hands), subjects could freely move their eyes.

Our naturalistic VR task is not only more intuitive to perform for all three species, it also allows us to capture behaviour in a much richer and more nuanced way than classical experimental paradigms. By analysing the paths the subjects take through the VR on each trial, we quantify their behaviour in a host of parameters beyond the classical reaction time (RT) and task accuracy (correct/incorrect). This allows us to quantify performance strategies and capacities in much more detail and on a single-trial level. For example, a high RT on a particular trial might indicate that an animal is strongly engaged in the task but unsure of the right answer (high level of attention), or that its focus is drifting off (low level of attention). Such differences can manifest in behavioural parameters such as the precision and efficiency with which stimuli are approached in the VR environment; as such, these parameters together give a more accurate manifestation of the attentional state of the subject. With this approach, we for the first time directly compare dynamics of spontaneous attentional fluctuations across three species, as well as the strategies they use to allocate fluctuating attentional resources to increase either response speed, accuracy or both - in a setting that is both naturalistic and highly replicable across species.

## Results

### A. Experimental Design

In order to compare attentional fluctuations in mice, monkeys, and humans, we engaged them in a simple two-alternative forced choice task, where they had to discriminate two leaf-shaped objects (Fig 1B). To elicit naturalistic fluctuations in attention, we employed an intuitive, foraging-based task structure (navigating towards food sources) based on naturalistic visual stimuli: leaves placed in a virtual reality (VR) environment consisting of a realistic grassy field and mountainous background, projected into a highly immersive dome completely surrounding the subjects (Fig 1A; see also (27)). The mice traversed this VR environment towards either object by running on a styrofoam ball; the monkeys and humans used a large trackball for this that they could operate with either one or both hands (monkeys), or the right hand only (humans). Subjects were freely viewing the stimuli and the environment.

**Fig. 1.**
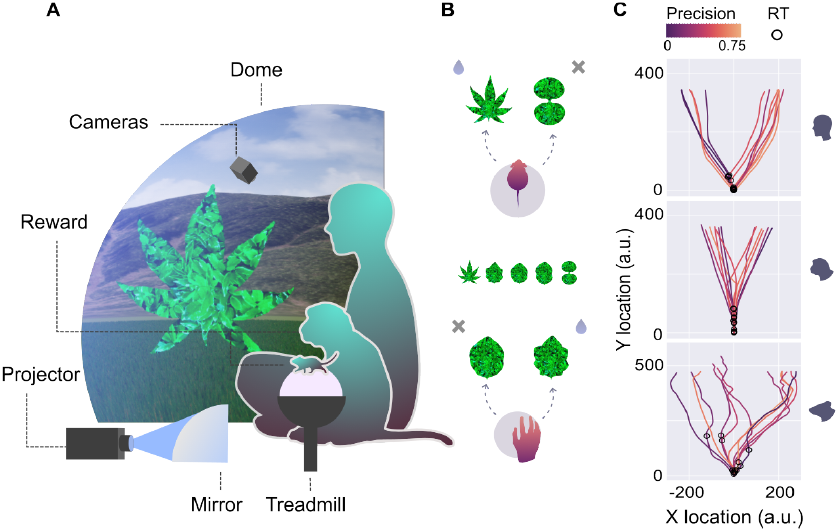
Experimental setup and path parameters. **A)** Mice, monkeys, and humans were placed inside a large dome, on which the VR task was projected via a curved mirror. Mice traversed the VR by running on a trackball, while monkeys and humans did so by moving a trackball with their hands. **B)** The perceptual decision task involved moving towards either of two stimuli, a rewarded target or an unrewarded distractor. **C)** The paths of the humans (top), monkeys (middle), and mice (bottom) towards the stimulus on several trials. Open circles denote reaction times (see Methods - Metrics for the definition) and path colour (on a normalised scale) denotes the ‘precision’ metric. Importantly, all metrics were normalised to each species’ average values; so even though the monkeys and humans had overall more precise paths than the mice, they also had more and less precise paths on individual trials, compared to their own average precision.

We used the subjects’ trajectories through the virtual environment to capture their task performance in five parameters, mapped onto the two seminal axes of attention: accuracy and speed. Capturing each axis by more than one behavioural parameter allowed us to make more nuanced estimates of speed and accuracy on a single-trial basis, beyond what can be achieved e.g. by binary classifications of hit and miss trials.

Three parameters specified task accuracy: *hit rate* (correct vs incorrect or miss trials), *precision* (how accurately either of the leaf-shaped objects is reached; see Fig 1C), and *bias* (the tendency to repeatedly approach objects on the same side, regardless of their identity). Both changes in precision and bias have previously been associated with different attentional states in mice (10, 13). The remaining two parameters specified speed: *reaction time* (the point in time the subject makes a turn towards one of the objects; see Fig 1C), and *speed* (average speed of moving through the VR environment from trial start to end). (For exact parameter definitions, see Methods - Metrics; for the distributions of these parameters for the three species, see Suppl Fig S1.)

### B. Models

The choice to categorize parameters as contributing either to the speed or accuracy dimension of task performance was made for conceptual reasons: we wanted to know how subjects navigated the trade-off between speed and accuracy given their available attentional processing resources. This trade-off has classically been considered a fundamental feature of attentional processing. To verify if this conceptually driven choice tallied with the actual structure of our behavioural data, we quantified the relations between the five behavioural parameters (Fig 2A, left). Specifically, we correlated the parameters’ overall time series (i.e. one value per trial, all sessions within one species concatenated). The resulting correlation matrix clearly showed two clusters of highly correlated parameters: those quantifying task accuracy and those quantifying speed, with low correlations between the clusters (Fig 2A, middle). This was the case for the correlation matrix averaged across all species, but also for the correlation matrices separately per species (Suppl Fig S2). As such, our data-driven analysis confirmed that behavioural parameters did indeed cluster onto the two axes of accuracy and speed, along which task performance could fluctuate independently.

**Fig. 2.**
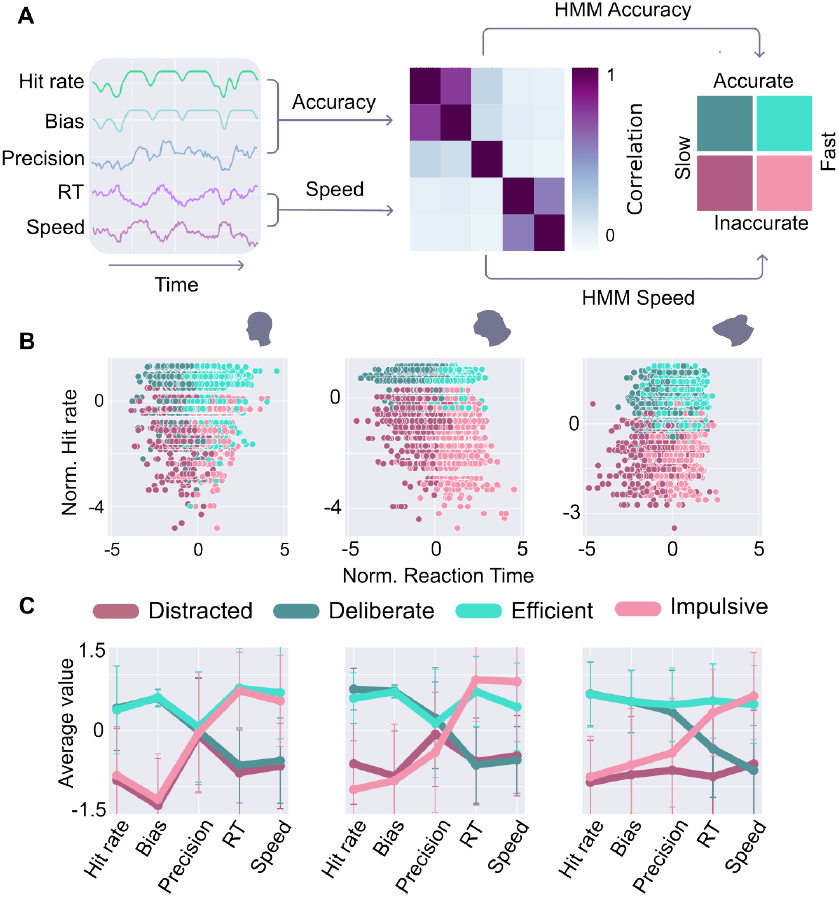
Attentional metrics and models. **A)** Five variables (left) were derived from the VR paths: three concerning task accuracy (hit rate, precision, and bias) and two concerning speed (running speed and reaction time). The correlation matrix of these variables (middle) confirmed that the variables concerning the classes accuracy and speed were highly correlated within each class, but largely independent between the two classes. The variables were therefore entered into two separate HMMs, the conjunction of which partitioned the data into four categories (right). **B)** Plotting a representative variable from each of the two classes (hit rate and reaction time) against each other and colouring the data by the four states, confirmed a correct partitioning of the data by the HMM conjunction, for humans (left), monkeys (middle), and mice (right). Legend of state colours: see C. **C)** The contribution of the five variables towards the partitioning into four states by the conjuncted HMMs was very similar between humans (left) and monkeys (middle). In mice (right), the precision variable had slightly different contributions towards the four states, but the overall pattern was similar to that of humans and monkeys. For this analysis, reaction time, bias and precision were inverted so that higher values in all metrics represent higher performance.

Attention is not the only factor shaping performance speed and accuracy - external factors like stimulus difficulty and history on a particular trial also play a decisive role. We hypothesised that dynamic changes in the two parameter clusters could reflect fluctuations in sustained attention beyond the performance differences elicited by specific trial attributes. To better capture such spontaneous fluctuations and minimize the contribution of external trial factors, we first smoothed the time series with a window size of 5 trials (see Methods, also Suppl Fig S4). To parcellate these smoothed time series into high and low attentional states, we used two independent Hidden Markov Models (HMMs; for details on the models, see Methods - HMMs). The accuracy and speed parameters were used as inputs into the two separate HMMs, each outputting two states: one HMM differentiating between slow and fast states, and one HMM differentiating between accurate and inaccurate states. The two HMMs were set up separately for each species; so even though humans and monkeys have generally better task performance than mice in absolute terms, relative task performance matters. The conjunction of the two HMM outputs resulted in four distinct state classifications: *distracted* (low accuracy and low speed), *deliberate* (high accuracy and low speed), *impulsive* (low accuracy and high speed), and *efficient* (high accuracy and high speed) (Fig 2A, right). Each task trial was assigned to one of these four states.

To verify that these four states are really present in the data and not artificially induced by our two HMMs, we coloured the trials according to the state they were assigned to and plotted the two parameters that are considered the most canonical measures of attention, hit rate and reaction time, against each other (Fig 2B). This analysis showed that indeed the conjuncted HMM states neatly partitioned the data into four quadrants, each with a different relationship between hit rate and reaction time. To check how well each of our five parameters partitioned the data as compared to the HMM states, we repeated this analysis with the remaining parameters (Suppl Fig S3). All parameters partitioned the data well, with the exception of precision in humans and monkeys, and reaction times in mice. We suspect this might be due to the humans and monkeys having extremely high precision on all trials, compared to the mice.

To complement the partitioning analysis, we investigated the relative representations of all five parameters in the two HMMs. As expected, in the accurate states, the three accuracy parameters had higher representation (and vice versa for the inaccurate states), whereas in the fast states, the speed parameters were represented strongly (Fig 2C). Importantly, this pattern was extremely similar across the three species, indicating that these input parameters, despite being recorded in very different ways (actual running for the mice, moving a trackball with the hand for monkeys and humans), are similarly important in determining the attentional state of the animal.

### C. State characterization

Classifying trials into the four attentional states and plotting them over time revealed that in all three species, subjects did not transition into new states constantly, but instead spent extended periods of time in one and the same state (see Fig 3A for an example session). The presence of such stable periods of the same state was not due to our smoothing procedure (Suppl Fig S4). Moreover, the overall proportion of time spent in any of the four states was quite similar between the three species (Fig 3B). Humans and monkeys spent comparatively more time in slow than in fast states; in humans, this was most pronounced (Fig 3B). Furthermore, more time was spent in accurate than in inaccurate states; in both monkeys and humans, this was due to a large proportion of distracted (slow and inaccurate) trials. Note that in the ‘distracted’ state, by far not all trials had slow and incorrect outcomes; just comparatively more than during the extremely good performance of the efficient (fast and correct) state.

**Fig. 3.**
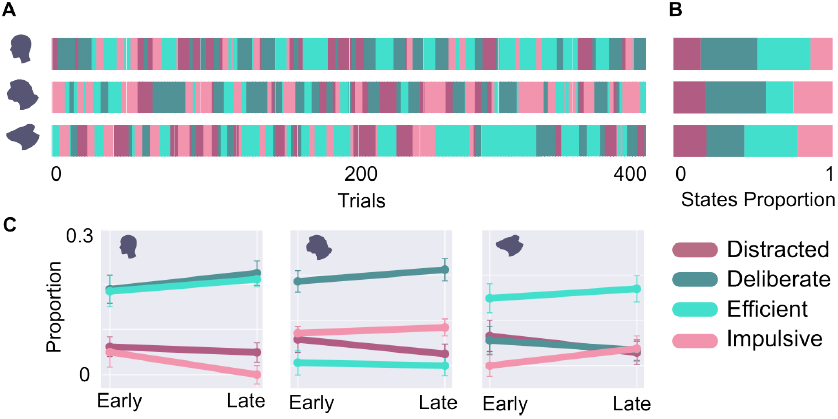
Model comparisons across species. **A)** Partitioning the trials of an example session of humans (top), monkeys (middle) and mice (bottom) into the four states showed both long (several tens of trials) and short (just a few trials) stretches of a particular state. **B)** Overall time spent in each state was very similar for humans and monkeys, whereas mice spent less time in the distracted, and more time in the deliberate state (histograms on the right). **C)** Distinguishing between early and late trials in the session did not yield any consistent pattern across species.

One potential confound is that performance speed and accuracy may be strongly shaped by internal variables that are not directly related to fluctuations in sustained attention, for instance, slow changes in fatigue (or in the case of monkeys and mice, satiety) throughout a session. While the drift correction implemented in our HMM analysis (see Methods, HMMs) should account for such slow changes to some extent, they may still bias our analysis throughout the time course of a session. To check whether slow drifts in internal variables were driving the occurrence of our four performance states, we computed the relative proportion of the four states over the course of each session by splitting sessions into an early and a late half. As shown in Fig 3C, the proportions of the four different states remained highly stable between the first and second half of the session for all three species (permutation tests yielded p-values > 0.05, see Methods). This suggests that the state transitions inferred here were not driven by slow internal drifts, and can thus be seen as a viable proxy for spontaneous attentional fluctuations.

### D. State dynamics

While the overall proportions at which the four different states occurred was largely comparable across species (see Fig 3B), their distribution over time may still vary between species. Anecdotally, many neuroscientists are familiar with the hugely different attention spans of the various species; mice can often only be trained for about 20-30 minutes at a time, whereas a typical human experiment lasts about an hour, and for a well-trained monkey, training times of up to 2-3 hours are not unusual. This suggests that state durations in mice might be shorter, and state transitions occur more often than in humans and monkeys. To test this hypothesis, we computed the dwell times for all states (Fig 4A). This showed that durations spent in each of the four states were on average roughly 6 trials, with significant but small differences of less than a trial between them (Kruskal-Wallis test, H = 14.8, p < 0.001). Slight differences in dwell times between the four states existed for each individual species. For example, humans spent comparatively longer uninterrupted sequences of trials in the accurate than in the inaccurate states (Fig 4A, left); this was not the case for monkeys (Fig 4A, middle) or mice (Fig 4A, right). Control analyses with smoothing over window sizes between 0 and 20 trials confirmed that dwell times scale linearly with window size, for all states and in all species; and therefore, any differences between species and states cannot have been artificially introduced in the data by smoothing (Suppl Fig S4).

**Fig. 4.**
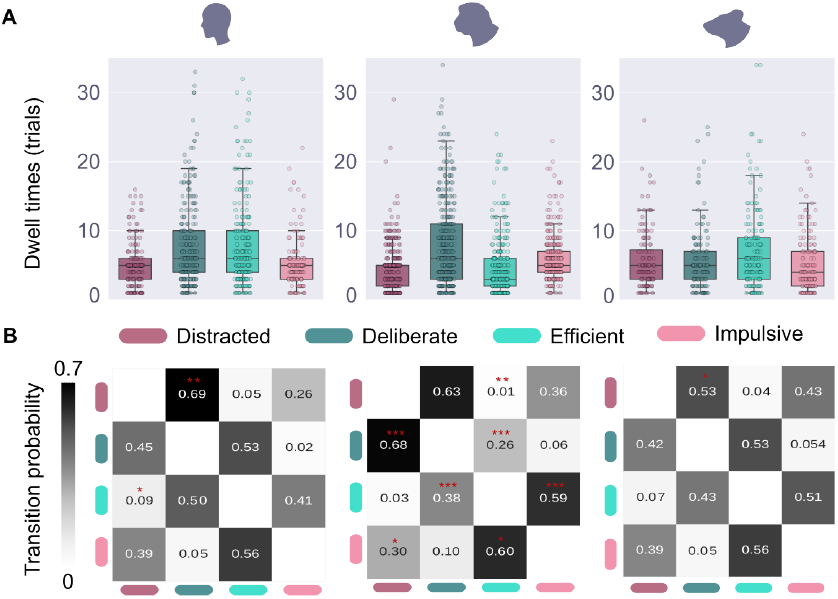
State temporal dynamics. **A)** Dwell times show a large spread, but are on average very similar between the species. Within each species, interesting differences emerged; for instance, humans, tended to stay longer within an accurate than within an inaccurate state. **B)** Transition matrices of humans (left), monkeys (middle), and mice (right) looked very similar. Transitions are more likely within a (speed or accuracy) class than between classes (’sketchy’ pattern), and for monkeys, transitions are more likely within the speed class than within the accuracy class (dark grey squares in top left and bottom right corner, compared to light grey bottom left - top right diagonal).

Another potential difference in the dynamic distribution of states over time is the sequence in which they tend to occur. For instance, as mice transition away from the efficient state, they might be more likely to move into an impulsive state, while monkeys or humans might be more likely to transition into a deliberate state. To explore whether there were specific sequences in which states were more prone to occur, we computed transition probability matrices, showing the probability of transitioning from any state to any other one (Fig 4B). The transition matrices for all three species looked strikingly similar. Most notably, a ‘patchy’ pattern emerged that implies much more likely transitions within one performance axis (accuracy or speed) than between axes. For example, a subject in the deliberate state (slow and accurate) is much more likely to switch to the distracted state (slow and inaccurate; so staying within the ‘speed’ axis), or to the impulsive state (fast and inaccurate; so staying within the ‘accuracy’ axis), than to the efficient state (fast and accurate; so switching both the speed and accuracy axis). In other words, for all three species, switching along both the speed and accuracy axis at the same time was highly unlikely. This could be an artifact of the simple base probabilities of a state switch coinciding across two independently computed axes. To test for this possibility, we computed histograms around each type of transition (c.f. peristimulus time histograms or PSTHs), which showed how often each of the states occurred around any other state. For all three species, transitions within the same (speed or accuracy) axis happened more often and transitions to the other axis happened less often than the baseline (Suppl Fig S5). Thus, the preference of staying along one axis was a real effect.

Examining the ‘within’ axis transitions more carefully revealed that in monkeys, transitions are much more likely within the ‘speed’ axis (for example, from deliberate to distracted, and from efficient to impulsive), than within the ‘accuracy’ axis (with the probability of a switch between deliberate and efficient states being significantly lower than shuffled data). This effect was highly significant (p>0.0005 in a two-tailed test with Bonferroni correction) and not present in humans or mice. We hypothesise that this might be due to the monkeys being extremely trained on the task; they might have been able to perform well even when distracted so that attentional fluctuations would manifest mostly in the speed with which they would perform the task.

Even though the dwell times show a large spread for each species and state (Fig 4A), it could be that there is a preferred dwell time, which would imply the existence of a certain rhythm in the state durations. To examine this, we determined the frequency spectrum of the time series of all trials in all sessions (Fig 5). As the transition matrices (Fig 4B) showed that switches between the speed and accuracy axes simultaneously were highly unlikely, we computed separate frequency spectra for each axis. Frequency spectra were computed using superlet analysis ((28); see also Methods - Frequency Analysis). To our surprise, the frequency spectra along the speed axis revealed a prominent, and highly significant peak for all three species (Fig 5A). This peak was at a period of about 55 trials in humans and monkeys; for mice, it was slightly slower, at a duration of about 70 trials. This suggests that there is an intrinsic, and potentially evolutionarily preserved, rhythm by which fast performance can typically be maintained for a period of tens of trials. In contrast, the frequency spectra of the accuracy axis did not reveal any significant peaks. Peaks that appeared most prominent were at a faster time scale of 20-40 trials.

**Fig. 5.**
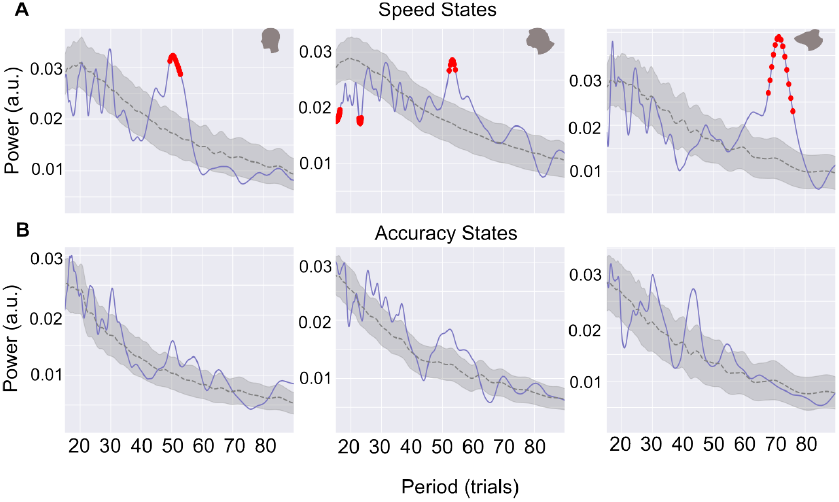
State frequencies. **A)** Frequency analysis on the speed states of humans (left), monkeys (middle) and mice (right) revealed a similar frequency peak in mice and monkeys around 55 trials, and a peak around 70 trials in mice (right). In all three species, frequency peaks were also seen between 10-30 trials, though these only reached significance in monkeys. Significance away from the mean (dotted line) is indicated in red. **B)** Same as A, but for accuracy states.

### E. Effect of external task attributes

The presence of a strong rhythmic component in the speed state fluctuations (and the absence of such a rhythm in the task structure) suggests that these fluctuations are internally generated rather than task-induced. To verify this, we made use of a task attribute that strongly affects task performance, namely task difficulty. For all three species, the task included trials of varying levels of difficulty (see Methods for details). We examined the effect of task difficulty by computing the average difficulty of the trials around the time of the state transitions (slow to fast states and fast to slow states). Any effect of task difficulty would have resulted in a peak or trough around the time of state transition; however, this was not observed (Fig 6). This suggests that the speed state transitions are entirely self-generated, despite the fact that more difficult trials typically lead to longer reaction times (29). Repeating this analysis for the accuracy states did show an effect of task difficulty (Suppl Fig S6). However, this effect is trivial because of the strong effect of task difficulty on task accuracy: a sequence of difficult trials will lead to increased incorrect performance, which is interpreted by the HMM as a transition from an accurate to an inaccurate state.

**Fig. 6.**
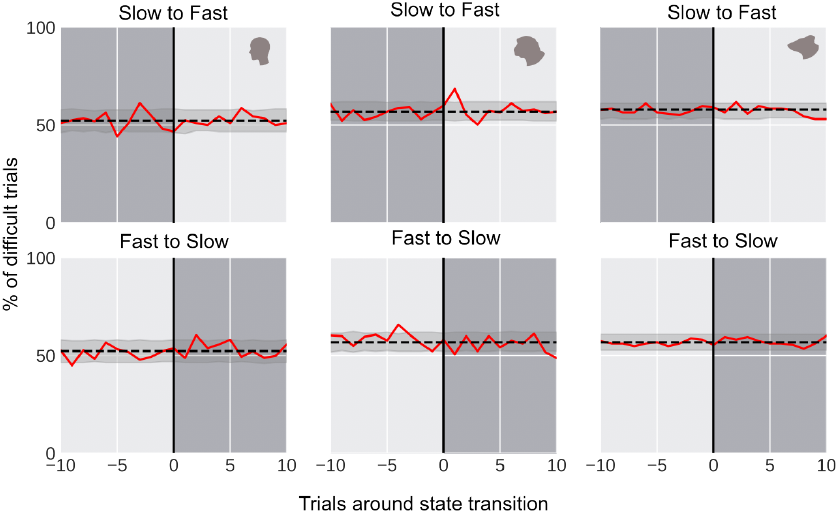
Effect of task difficulty on state transitions. Around the time of a state transition (vertical line) from slow to fast (top row) or fast to slow (bottom row), the difficulty level of trials (red line) was not significantly different from the average difficulty (dotted line). The grey band shows the range of 1000 shuffled controls (see Methods). This indicates that the transitions between speed states are internally generated, as opposed to externally triggered, in all three species.

## Discussion

Sustained attention is an important cognitive process underlying the ability of flexible behaviour in a diverse range of mammalian species (30, 31). Yet the way it has been studied to date has been very species-specific, precluding meaningful cross-species comparisons. Here, we engaged mice, monkeys, and humans in the exact same, naturalistic perceptual decision task in a VR environment. We captured their behaviour in five parameters that were based on their paths through the VR, and that described their task performance either in terms of task accuracy or speed, two seminal task-dependent variables modulated by attention (11, 12). Speed and accuracy parameters served as inputs into two separate HMMs to infer underlying attentional states. The conjunction between these two separate HMMs partitioned the trials for all three species into four attentional states: a distracted, deliberate, efficient, and impulsive state.

We found that when attention was defined relative to the overall capacity of the animal for the task at hand, all three species were immersed in the four states for roughly the same amount of time (Fig 3B). Moreover, they displayed the same overall behaviour; short stretches (6 trials on average, see Fig 4A) of being in the same state, with frequent transitions between them (Fig 3A). These transitions preferentially happened either along the speed or accuracy axis; transitions from the slow and inaccurate state to the fast and accurate state or vice versa were rarer than predicted by chance (Fig 4B). Although the short stretches of staying in the same state could vary widely in duration (Fig 4A), a cycle of a slow and a fast state lasted preferentially for around 55 trials for humans and monkeys, and for around 70 trials for mice (Fig 5A). Transitions between accuracy states did not reveal any significant cycles. This rhythm suggests that the attentional fluctuations are internally generated; the absence of an effect of task difficulty on the transition confirms this (Fig 6). Summarising, fluctuations in attention were remarkably similar between the species.

HMMs, often in combination with a general linear model (GLM-HMMs) to account for the underlying states dependence on continuously incoming sensory information (32), have been used before to delineate states of high and low sustained attention. For example, in mice involved in perceptual decision-making tasks, a GLM-HMM trained on task accuracy distinguished between either one engaged and two disengaged states (15), or between an optimal, sub-optimal, and disengaged performance state (10). In fact, when we trained a more recent type of HMM, called a Markov Switching Linear Regression (MSLR) model, on the reaction times in the same visual discrimination task as employed here, we revealed underlying internal states that mapped onto differences in reaction time and task accuracy in both mice and monkeys (33).

It is worth noting that the attentional states described in these studies typically lasted much longer than the states in our data, on the order of tens up to roughly a hundred trials (c.f. example data in (15), Fig 1F, and (10), Fig 1E). A possible explanation for this might be found in the fact that we enforced two states onto the data, whereas the GLM-HMMs described above all had at least three states; in mice involved in a perceptual decision-making task, a two-state GLM-HMM resulted in single trial switches, whereas a three-state GLM-HMM on the same data set yielded much slower transitions (34). Also, overall switch rate (but not relative switching proportions) depend critically on the smoothing parameters chosen (see Suppl Fig S4). Lastly, it might be possible that our task simply induces faster switching between states; the MSLR model trained on the same data yielded three states for mice and four state for monkeys, yet showed similarly short dwell times as shown in the current analysis (33).

Overall, we found that attentional dynamics in all three species were very similar. Attentional state durations were similar, even including a strikingly similar peak in the frequency spectrum, as well as the transitions between them. These measurements were reported in the time unit of trials; reporting the same results in seconds would have slightly altered these outcomes owing to the higher average trial duration for the mice (see Suppl Fig S7). However, it is very likely that trials are longer in mice due to a different mode of moving through the VR (which is independent from the task per se); and since all our measurements are done at trial resolution (i.e. we do not distinguish between various events within a trial), we believe that reporting our results in trials is the most accurate depiction of the attentional fluctuations.

The similarity in dynamics between the species might be expected based on the supposedly common underlying neural mechanisms of attention in primates and rodents (35, 36). Similar perceptual decision-making tasks for different species have revealed subtle differences between the species. For example, in a new version of the pulse-based evidence accumulation task (in which subjects have to select the right or left side where light pulses are flashed with the highest probability), it was shown that humans prioritize accuracy, whereas rodents favour speed of responding (37). This is in line with our finding that mice were in a fast (efficient or impulsive) state a larger part of the time than humans and monkeys. And in an uncertain decision-making task for mice, monkeys, humans, which was designed to investigate the balance between exploiting one strategy to solve the task and exploring alternative strategies, mice were less persistent in exploitation behaviour than primates (38). The development of tasks such as these is crucial but difficult; even a simple component of a task such as the same food reward across species can have different value for these species (39).

In the monkeys, the probability of a transition between accuracy states was significantly lower than in mice and humans. One reason for this could be overtraining, i.e. the building of habitual responses, which given the long training times for the monkeys, might have affected them. Research on habit learning suggests that what initially starts out as goal-oriented behaviour often becomes habitual with extensive practice (40). The relationship of attention with habitual behaviour is complex - heightened attention can trigger habitual responses (41), and habitual responses can take over even if attention is directed elsewhere (42). Therefore, it could be that, even though attention wandered, the monkeys still performed with high accuracy, and these attentional fluctuations mainly manifested themselves within the speed states.

Despite the possible impact of overtraining, we believe our task was natural enough to elicit natural fluctuations in sustained attention. The fact that the attentional fluctuations seem not to be induced by task difficulty, but rather entirely self-generated, confirms this. An experimental set-up in which the animals and humans could freely move would have allowed for natural indicators of alertness, such as ‘head scanning’ movements in rats (43), and general body and head direction in primates (44). Even so, it is important to emphasise that VR set-ups (including the one we employed) are one of the only ways to keep tasks and visual input identical between different species, and are therefore ideally suited to cross-species comparisons (45). Moreover, our VR set-up allowed us to capture behaviour in exactly the same five parameters across the three species, which guarantees a direct and meaningful comparison between them.

We believe that this approach has value beyond just characterizing the common dynamics of sustained attention across species. A staggering 96 percent of all neuropsychiatric clinical trials fail (46). To successfully translate results from preclinical work to the clinic, comparable and sensitive tests of cognitive processing in animals are essential (25). However, such tests are difficult to conceive, as changes in behaviour as a result of neurological or psychiatric conditions are often difficult to reproduce in animals (47–49). An intuitive task such as the one presented here can help refine animal models of e.g. ADHD (50) and capture behavioural changes resulting from clinical interventions in much more detail. As such, we believe that developing tools to elicit and precisely quantify naturalistic behaviours across species in a directly comparable way is a crucial step in designing truly translatable cross-species tests of neuronal processing.

## ACKNOWLEDGEMENTS

We thank Gaby Schneider for useful input on the study. M.G. acknowledges support from the Fazit Stiftung.

## Methods

### F. Subjects

Three mice (FVB/BL6, male, 10-25 weeks old), two monkeys (Macaca mulatta, both male, 15 years old), and 11 humans (3 male, 8 female, mean age 25 years old) were used in this study. All animal experimental procedures were approved by the local ethics committee, Regierungspraesidium Darmstadt, under number F149/2000. The human experiments were approved by the ethics committee from the medical department of the Goethe University in Frankfurt under number 2021-252.

### G. Experimental Setup

Details of our experimental setup have been published previously (27). Briefly, participants were placed inside a spherical dome (diameter 120 cm, 250°) onto which a virtual reality environment was projected via a curved mirror. The virtual reality was created using the game engine Unreal 4 and consisted of a grassy landscape with mountains in the background and a blue sky overhead, in the middle of which the two stimuli were displayed. In order to navigate the environment, macaques and human participants used a GK75-1602B 75mm trackball from NSI. Mice were placed on top of a 20 cm diameter Styrofoam ball suspended in the air (modified method from (51)). The movements of either device were translated into movement in the virtual environment.

### H. Task design

All three species performed the same simple two-choice visual discrimination task. They were presented with two stimuli with naturalistic shape and texture (see Fig 1B). The target resembled a pointed-lobed leaf; the distractor resembled an hourglass. The perceptual similarity between target and distractor was varied systematically by morphing the shapes into each other. Both shapes were filled with a uniform foliage texture in a blue-green colour. The colour was selected to be easily visible both for rodents and primates. The participants were required to navigate towards and then (virtually) collide with the rewarded stimulus while ignoring the distractor.

The task had several difficulty levels; on easy trials, subjects had to distinguish between two shapes clearly resembling a pointed leaf and an hourglass, whereas on difficult trials, they distinguished between two morphed stimuli only slightly resembling a pointed leaf and an hourglass. Monkeys and humans were shown shapes of comparably high difficulty. Mice were presented with more discriminable stimuli, as the goal was not to test the limits of their visual discrimination ability, but to produce a reliable behavioural response.

### I. Metrics

Each trial could have one of three outcomes: correct response (collided with the target stimulus), incorrect response (collided with the distractor) and miss (no stimulus chosen). On each trial, the path of the participant through the virtual environment was recorded and used to extract five metrics. *Hit rate* was classified as a binary metric (correct (1) vs incorrect or miss (0)) and then smoothed over by the smoothing window (see Methods - Data Preprocessing). *Precision* was defined as the deviation (VR units) of a stimulus collision point from the average target collision point on the left or on the right side. *Bias* marked trials in which the participant chose the stimulus on the same side at least twice in a row, while the target stimulus position changed. *Reaction Time (RT)* was defined as the first significant change in direction in the path towards one of the stimuli (see Suppl Methods). Finally, *speed* refers to the average movement speed (VR units/s) of the participant in each trial.

### J. Data preprocessing

The data was preprocessed in several steps. First, we performed detrending on the time series of reaction time and speed by fitting a second-degree polynomial. This was necessary to remove the motor learning effect in some sessions, especially in naive human participants. The gradual increase in speed and decrease in reaction times levelled out after approximately 150-200 trials. The detrending step preserved the short-term fluctuations in the data while removing the slow trend where it was present. In the next step, we smoothed the data by applying a centred rolling average with a window of 5 trials to reduce the effect of single-trial difficulty. Finally, we applied the standard scaler to normalise the data. Within one species, each session was preprocessed individually and concatenated for further analysis.

### K. HMMs

First, the five metrics were correlated with each other. Based on the resulting correlation analysis, they were divided into two axes: accuracy (hit rate, precision and bias) and speed (reaction time and speed). These two axes provided the input to two hidden Markov models (HMMs). For this analysis we used an HMM with Gaussian observation class, implemented in an SSM package (52). As this is a hypothesis-driven analysis, we chose to split the data into 2 states. We also used sticky transitions with a stickiness parameter 50, based on the previous analysis (33). The models were trained on all the sessions concatenated, separately for each species. In order to offset state probabilities around the end of one session and the beginning of the next, we added 50 trial padding with a single value across metrics. The padding was reliably classified as a third state and discarded afterwards. Each model classified the trials as belonging to either of two states (accurate / inaccurate for the accuracy model, and slow / fast for the speed model). The conjunction of these classifications resulted in four states, occupying the quadrants of the speed/accuracy space: distracted (slow and inaccurate), deliberate (slow and accurate), efficient (fast and accurate), and impulsive (fast and inaccurate). Thus, each trial was assigned to one of these four states.

### L. Shuffling

We used shuffling to assess the significance of observed behavioural patterns. To create a null distribution we randomly shuffled the behavioural values of the trials a thousand times. The shuffled trials belonged to the same recording session and difficulty level (morphing stage - see Task Design). We then applied the same data transformations that we used on the original data set, including detrending, smoothing and scaling. Following that we trained the HMMs and compared the outcomes of the shuffling to the outcomes of the real experiment. If the relevant behavioural value of the real data was lower than 2.5 percent or higher than 97.5 percent of the average shuffled value, it was considered significant.

### M. Analysis of dwell times and transitions

Blocks of consecutive trials assigned to the same state were identified, and the times that such blocks lasted were termed the dwell times. Dwell times were statistically compared between the species using a Mann-Whitney U-test. Subsequently, transitions between the four states were computed and summarised in a transition matrix for each species. These matrices show how likely a trial of one of the four attentional states is followed by a trial of any (other or same) attentional state. To get a clearer picture of the transitions between states we subtracted the diagonal of the transition matrix, effectively excluding the self-transitions.

### N. Frequency analysis

In order to identify any regular fluctuation patterns we applied ‘superlets’ to the data. ‘Superlets’ are a tool, similar to wavelet transform, used to analyse the frequency content of a signal. Unlike other frequency analysis techniques, ‘superlets’ allow for ‘super-resolution’ in both time and frequency domains, making it possible to identify short bursts of oscillations (28). In order to identify prominent periodic patterns in behavioural data we focused on states of speed and states of accuracy separately. This way we could decompose the binary signal (states 1 and 0) into the constituent frequencies. The states labels of all the sessions from a single species were concatenated and then padded with a mirror-reflected signal. In the case of our data the time unit is one trial. We set the sampling rate to 1000 for convenience. We analysed the frequencies in the range of 6.7 (cycles/1000 trials) to 66.7 (cycles/1000 trials). This is equal to a range of 150 to 15 trials per cycle. The outcome of this is frequency power over time for each of the sessions. We then computed an average power value per frequency, averaged across sessions. This resulted in a power spectrum with several peaks.

We shuffled the raw data in a way that preserves original task structure but disrupts any potential slow behavioural effects, like attention. We randomly shuffled trials 1000 times within each session and within the perceptual difficulty level (only shuffling trials with the same stimulus morphing percentage). We then applied the same pre-processing steps on each of the surrogate datasets as we did on the original dataset. This includes smoothing with a 5-trial centred moving average window and scaling the data by computing z-score values. Following that we applied the HMM analysis and the ‘superlets’ analysis described earlier to each of the shuffled datasets.

Finally we computed the average power per frequency of each of the shuffled datasets and used these power spectra as a null-distribution to compare to the original dataset. We z-scored the original dataset using the mean and the standard deviation of the null-distribution and used 5th and 95th percentiles of the surrogate distribution as the significance thresholds. Values in the original data exceeding these thresholds were considered significant and highlighted in Fig 5.

## Supplementary Methods

### Reaction Time

In our VR setting, where the perceptual decision task entails moving towards one of two stimuli, reaction times (RTs) are defined as the time point of the initial substantial movement directed towards either stimulus, irrespective of minor positional adjustments. To calculate the RT, we use a sliding window linear regression approach, incorporating a time decay mechanism. This approach has been described in detail elsewhere (33) and will only be repeated briefly here.

First we computed a linear regression on the time series of lateral VR movement for adjacent sliding windows. Then, we added the time decay by predicting the movement for the window immediately following the window of interest and comparing it to the actual movement in that window. This gave us an array of movement values. From this array, we detected the local maxima. Once we found those, we further required that they have a minimum prominence. Prominence is a measure of the significance of a peak by comparing the peak to its surroundings. For the sake of stability, we used multiple window sizes (100, 150, 200 and 250 ms) to detect the lateral movement peaks and their prominence, and combined the results. The time of the largest movement peak was then taken as the RT.

## Supplementary figures

**Fig. S1.**
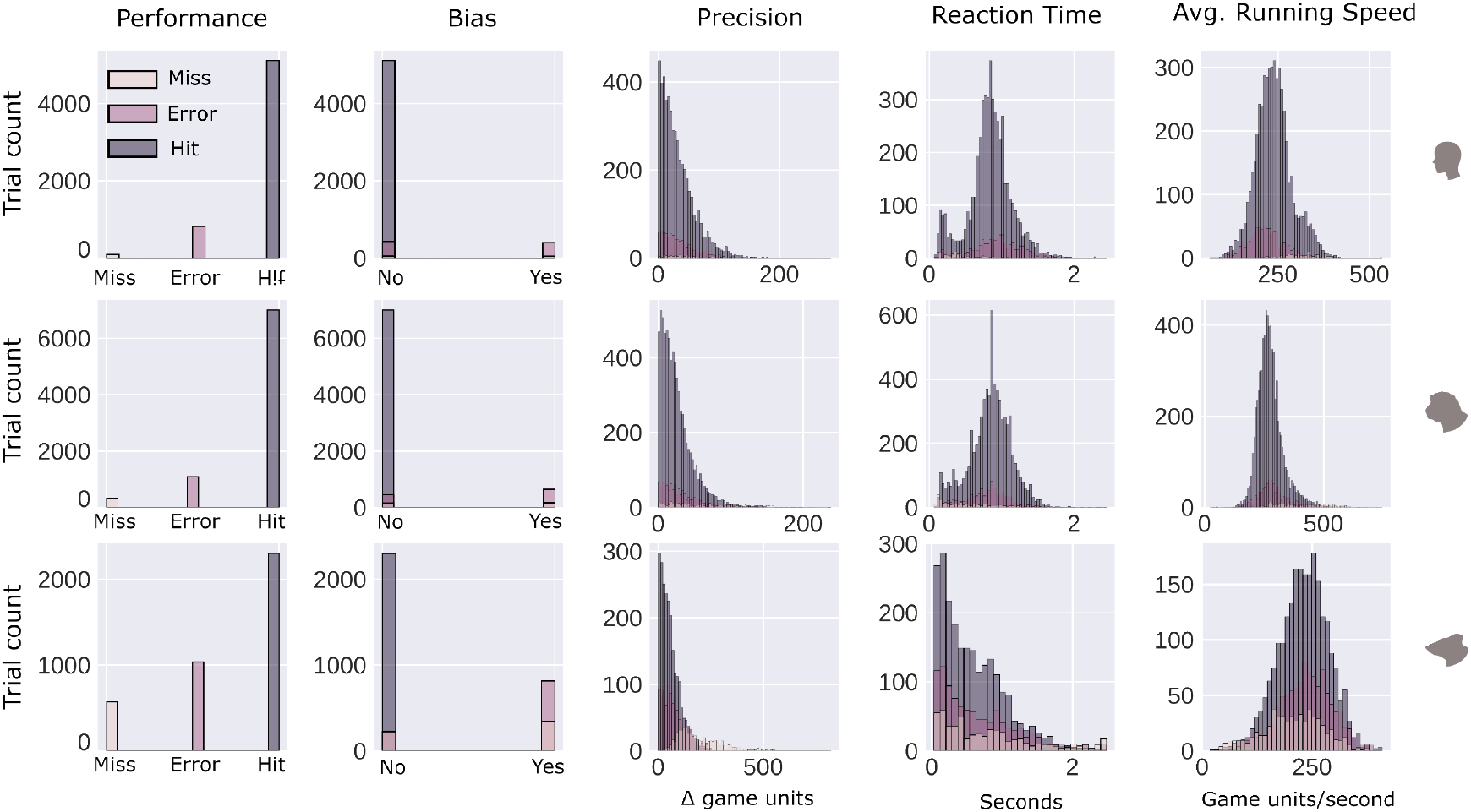
Input parameters for each species. Distributions of the five input parameters into the two HMMs are shown for humans (top), monkeys (middle) and mice (bottom).

**Fig. S2.**
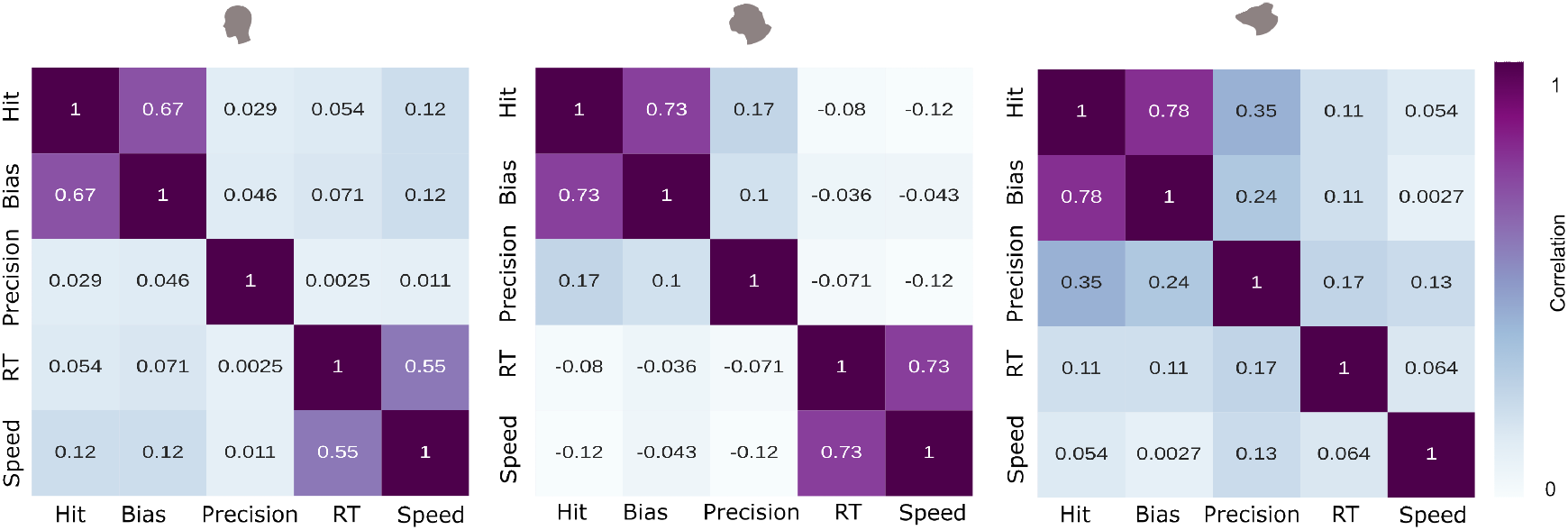
Cross-correlation matrices of behavioural parameters. The correlations between all parameters are shown for the three species separately; humans (left), monkeys (middle), and mice (right).

**Fig. S3.**
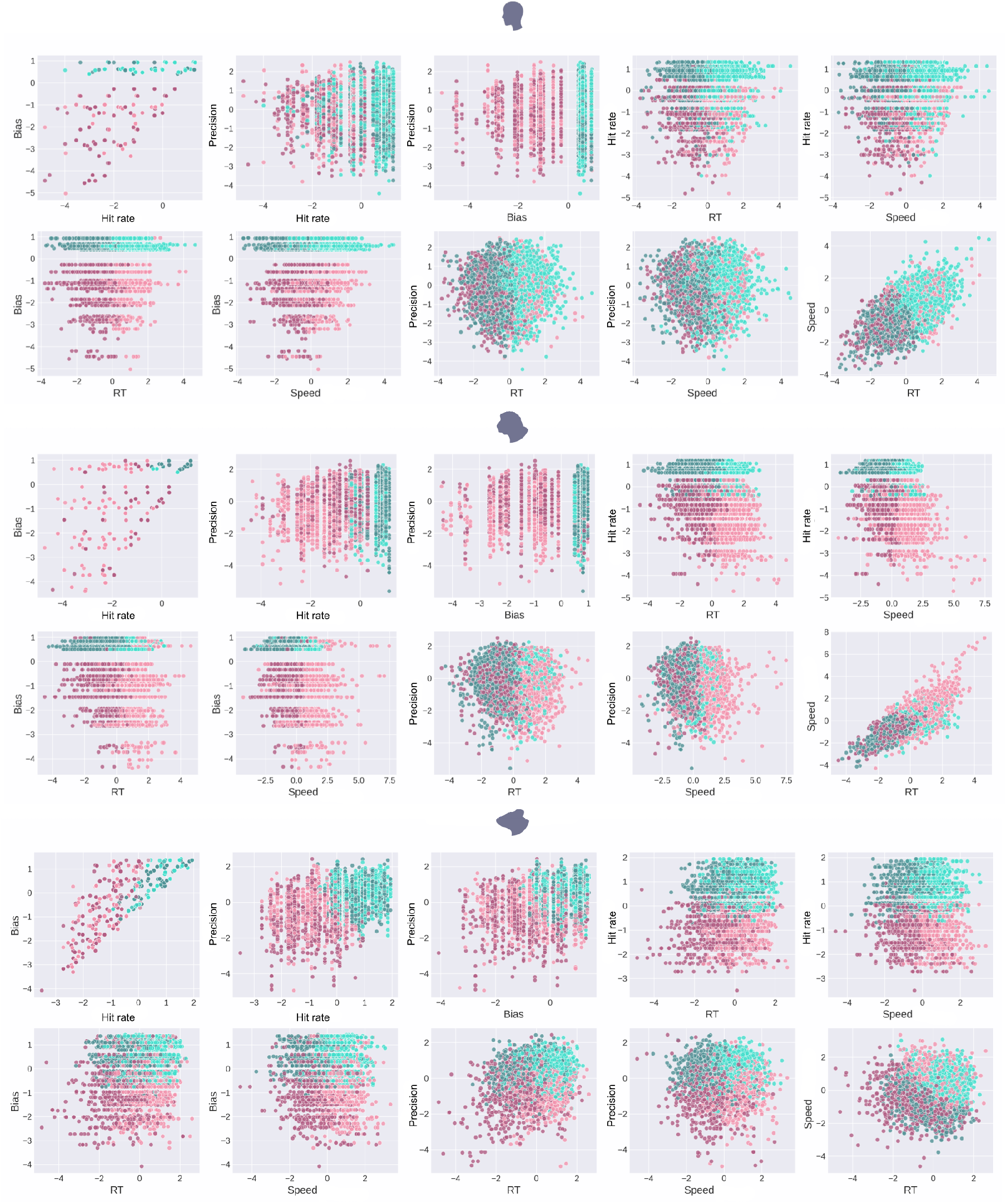
Comparison of individual parameters to HMM states. Plotting all variables against each other and colouring the data by the four states confirms a correct partitioning of the data by the HMM conjunction, for humans (top), monkeys (middle), and mice (bottom).

**Fig. S4.**
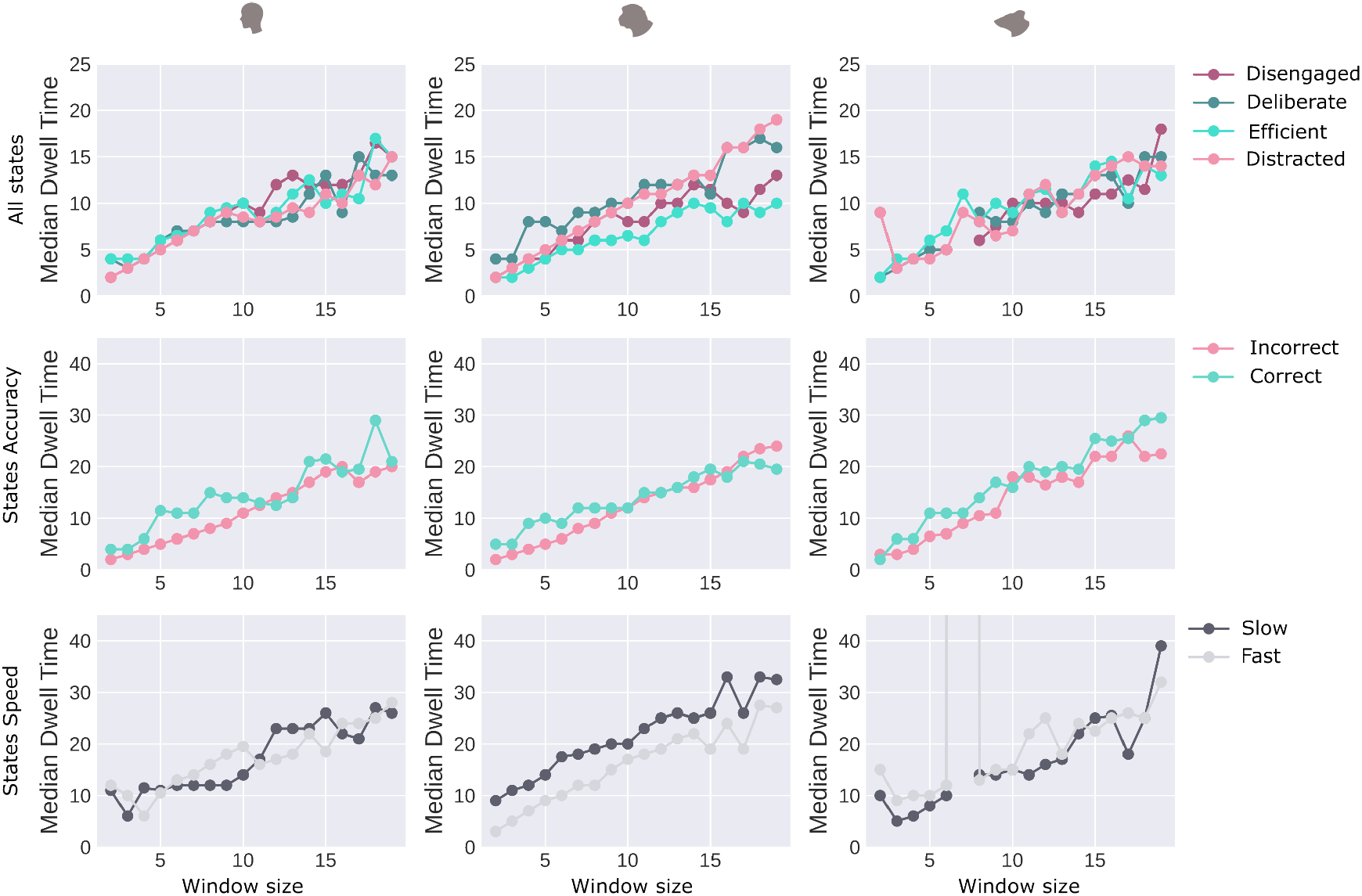
Effects of smoothing. Increasing the window size linearly increased the dwell times for all states and in all species; thus, window size has no effect on the relative dwell times between the states.

**Fig. S5.**
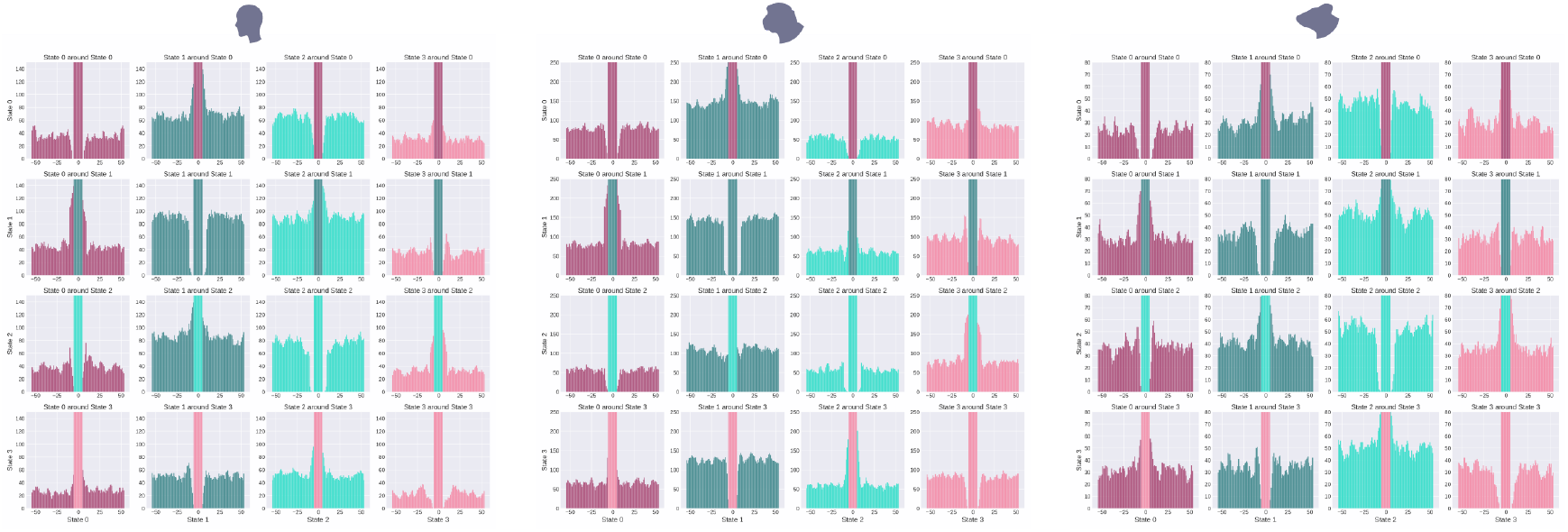
Occurrences of state transitions. In all three species, for each state, occurrence of a transition to any other state in the time immediately preceding and following a transition into that state. Certain state transitions happen more often (e.g. between 0 and 1), while others happen less often (e.g. between 0 and 2).

**Fig. S6.**
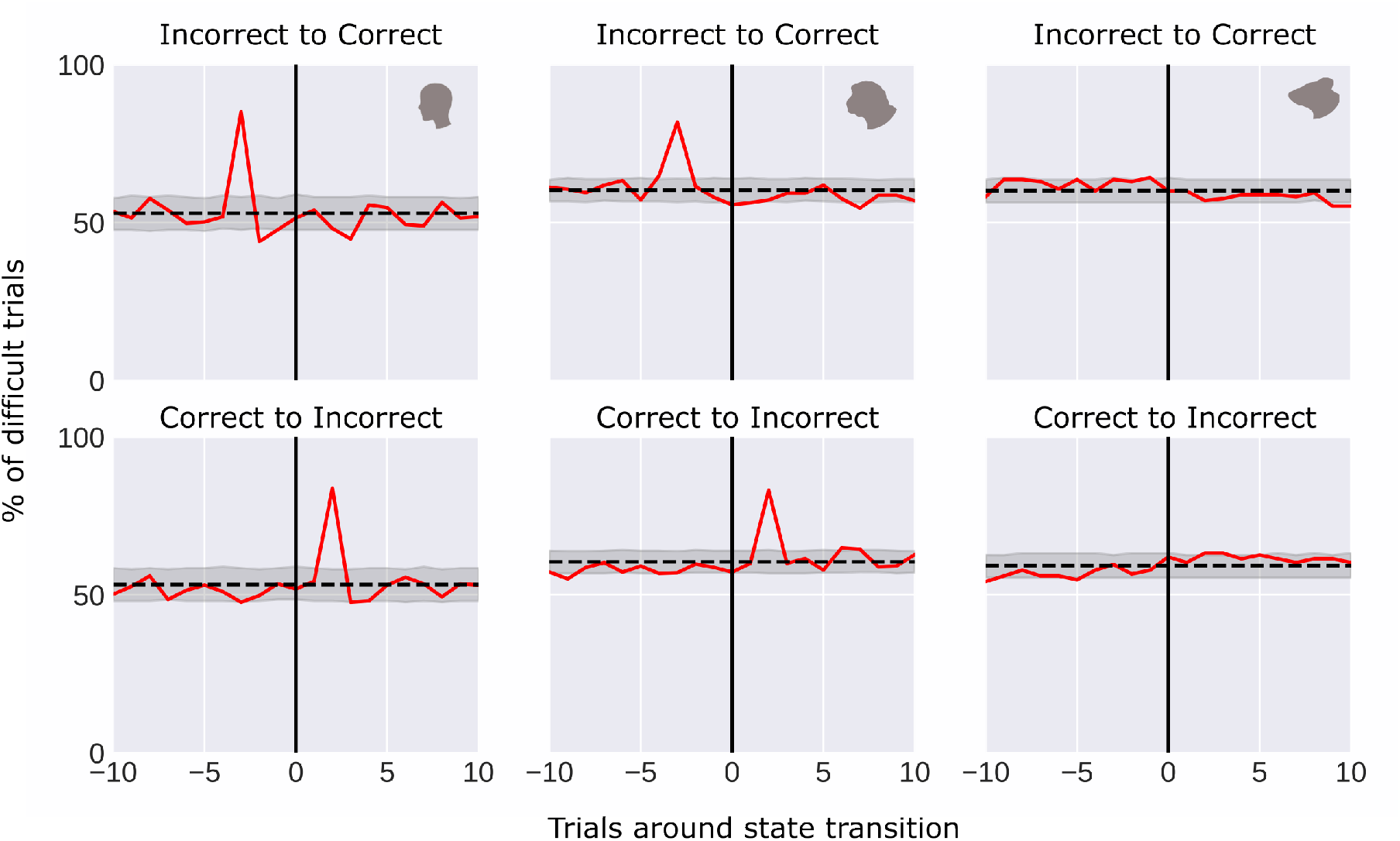
Effect of task difficulty on accuracy state transitions. A difficult trial will cause a transition from correct to incorrect in monkeys and humans. The reverse effect is also present.

**Fig. S7.**
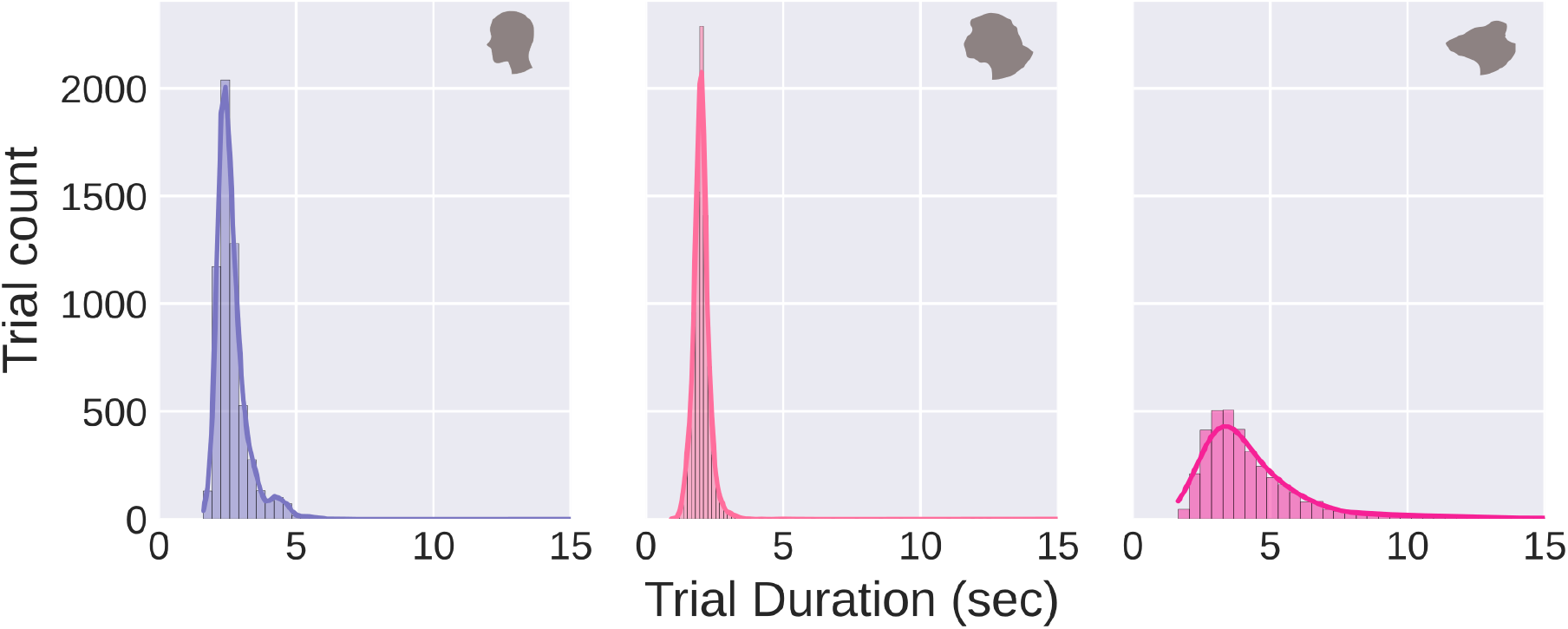
Trial durations. Trials durations in seconds, for the three species. Trial durations are similar between humans and monkeys, and longer in mice.

